# Obesity influences composition of salivary and fecal microbiota and impacts the interactions between bacterial taxa

**DOI:** 10.1101/2021.10.12.464168

**Authors:** Andrei Bombin, Jonathan D. Mosley, Shun Yan, Sergei Bombin, Jane F. Ferguson

**Affiliations:** Division of Clinical Pharmacology, Department of Medicine, Vanderbilt University Medical Center, Nashville TN; Department of Biomedical Informatics, Vanderbilt University Medical Center, Nashville TN; Department of Genetics, The University of Alabama, Birmingham AL; Department of Biological Sciences, The University of Alabama, Tuscaloosa AL; Division of Cardiovascular Medicine, Department of Medicine, Vanderbilt University Medical Center, Nashville TN

## Abstract

Obesity is an increasing global health concern and is associated with a broad range of morbidities. The gut microbiota are increasingly recognized as important contributors to obesity and cardiometabolic health. This study aimed to characterize oral and gut microbial communities, and evaluate host:microbiota interactions between clinical obesity classifications. We performed 16S rDNA sequencing on fecal and salivary samples, global metabolomics profiling on plasma and stool samples, and dietary profiling in 135 healthy individuals. We grouped individuals by obesity status, based on body mass index (BMI), including lean (BMI 18-24.9), overweight (BMI 25-29.9), or obese (BMI ≥30). We analyzed differences in microbiome composition, community inter-relationships, and predicted microbial function by obesity status. We found that salivary bacterial communities of lean and obese individuals were compositionally and phylogenetically distinct. An increase in obesity status was positively associated with strong correlations between bacterial taxa, particularly with bacterial groups implicated in metabolic disorders including *Fretibacterium*, and *Tannerella*. Consumption of sweeteners, especially xylitol, significantly influenced compositional and phylogenetic diversities of salivary and fecal bacterial communities. In addition, obesity groups exhibited differences in predicted bacterial metabolic activity, which was correlated with host’s metabolite concentrations. Overall, obesity was associated with distinct changes in bacterial community dynamics, particularly in saliva. Consideration of microbiome community structure, and inclusion of salivary samples may improve our ability to understand pathways linking microbiota to obesity and cardiometabolic disease.

**IMPORTANCE:** Obesity is a worldwide epidemic that is associated with a wide range of health issues. Microbiota were shown to influence metabolism and obesity development. Our study aimed to evaluate the interactions between obesity, salivary and fecal microbiota, and metabolite concentrations in healthy individuals. The oral bacterial community was more impacted by the obesity status of the host than fecal microbiota. Consistently for oral and fecal microbiota, the number of strong interactions between bacteria increased with the increase in the obesity status. Several predicted microbial metabolic pathways that were shown to be associated with metabolic health were uniquely enriched between obesity groups. In addition, these metabolic pathways were correlated with plasma and stool metabolites. Our results suggest that oral microbiota might better reflect the obesity status of the host than fecal microbiota, and that correlations between microbial taxa are altered during obesity.

## INTRODUCTION

Obesity is a growing worldwide epidemic and is linked to a range of health issues including hypertension, type 2 diabetes, asthma, coronary heart disease, Alzheimer’s disease and cancer [1-5]. Known risk factors include imbalances between calorie intake and expenditure, genetics, stress, and disruptions in the endocrine system [1, 6]; however much remains unknown. Better characterization of mechanisms predisposing to obesity could enable novel prevention and treatment strategies.

The composition of an individual’s microbiota is increasingly being recognized as a contributor to obesity risk [7-9]. Microbiota can influence the host’s metabolic phenotype both by directly affecting energy and nutrient availability [10-14], and through modulation of signaling pathways [15-22]. Previous studies suggested that the fecal symbiotic bacterial community of obese individuals is less diverse than that of lean individuals [8, 23]. In addition, the abundance of several bacterial taxa including *Lactobacillus, Pervotella, Alistipes, Akkermansia*, and others vary with obesity status [7, 9]. Salivary microbiota of lean and obese individuals also differ in diversity and composition [9, 24-27]. Abundance of several salivary bacterial taxa including *Campylobacter, Aggregatibacter*, and *Veillonella* was reported to be positively associated with obesity [28-30]. Higher abundances of Bacteroidetes, Spirochaetes, and Firmicutes were observed in lean individuals [9, 31, 32]. However, data are contradictory, even for rather abundant bacteria taxa. For example, the abundance of intestinal *Lactobacillus* was reported to be both positively and negatively associated with obesity [7, 33-35]. These discrepancies may be due in part to complex interactions between microbial community members, where metabolic activity of individual bacterial taxa can vary based on the activity of other microbes in the community [36-39]. Consideration of interactions between members of microbiota might be essential to improve identification of bacterial mechanisms underlying obesity.

We hypothesized that the presence of obesity, in the absence of known disease, would associate with differences in microbiome composition and function. We further hypothesized that community structure and bacterial inter-relationships would differ by obesity status. We evaluated the differences in compositional and phylogenetic diversity of salivary and fecal microbiota between obesity groups in a well-characterized sample of healthy individuals. We examined interractions between bacterial taxa based on obesity status of the host, and showed that predicted bacterial metabolic activity varies between obesity groups and is correlated with intestinal and circulating metabolite concentrations.

## MATERIALS AND METHODS

### Study population

We analyzed data from the ABO Study (n=135) as described previously [40-42]. Briefly, healthy non-pregnant and non-lactating women and men were recruited to a cross-sectional study. Participants completed dietary profiling (validated 3-day food records, and DHQ II food frequency questionnaires [FFQ]), and provided stool, saliva, and blood samples. Height and weight were measured at the study visit. Individuals were classified based on body mass index (BMI, weight (kg)/height (m)-squared), including lean (BMI 18-24.9; fecal samples n=76, saliva samples n=49), overweight (BMI 25-29.9; fecal samples n=34, saliva samples n=19), or obese (BMI ≥30, fecal samples n=25, saliva samples n=16), to explore differences in composition and function of microbiota by obesity. All participants provided written informed consent. The study was approved by the Institutional Review Boards of the University of Pennsylvania and Vanderbilt University.

### Sample Profiling

As we have previously described, 16S rDNA sequencing of the bacterial V4 fragment was performed on Illumina MiSeq platform using 135 fecal and 85 saliva samples to identify bacterial community composition [42]. Global metabolomics profiling of fecal and plasma samples, from a subset of individuals (n=75) was performed at Metabolon (Metabolon Inc., Morrisville, NC, United States).

### Pre-analysis processing

#### Sequences alignment and normalization

Pre-analysis processing of 16SrRNA reads was performed with R v4.0.2 [43]. Demultiplexed sequences were filtered, forward and reverse reads were merged, and resulted sequences were assigned to amplicon sequence variants (ASVs), with the default settings of DADA2 pipeline v1.18.0 [44]. Chimeric sequences were also removed with the dada2 package v1.18.0 [44]. Sequence variants were assigned taxonomy with dada2 and SILVA v138.1 database [44, 45]. ASVs counts were normalized with cumulative sum scaling method implemented in the metagenomeSeq v1.32.0 package [46]. In the salivary samples, we identified 1,932 ASVs that belonged to 12 phyla, 19 classes, 44 orders, 70 families, 134 genera, and 229 bacterial species. In our fecal samples, we identified 5,000 ASVs that belonged to 16 phyla, 26 classes, 55 orders, 86 families, 270 genera, and 338 bacterial species.

#### Alpha diversity

Normalized ASVs counts were used to calculate species richness, Shannon, and Gini–Simpson alpha diversity indices with the vegan v2.5.7 package [47]. <u>Beta diversity.</u> Bray-Curtis distances were calculated with vegan v2.5.7 [47]. Unrooted neighbor-joining tree was computed with the ape package v5.5 [48]. The tree was optimized based on generalized time-reversible model implemented in the phangorn v2.5.5 package [49, 50]. Lastly, weighted and unweighted Unifrac distances between each sample were calculated with the phyloseq v1.30.0 package [51].

<u>Functional potential</u> of the bacterial communities was predicted with PICRUSt2 according with the default pipeline [52]. Predictions were made for Enzyme Commission numbers (EC), Kyoto Encyclopedia of Genes and Genomes orthologs (KO), and MetaCyc pathways [52-55]. In accordance with PICRUSt2 authors’ recommendations, the resulting data were transformed with centered-log ratio transformation implemented in the ALDEx2 v1.24.0 package [56].

### Statistical Analysis

Statistical analysis and data visualization was done with R v3.6.1 [43]. Beta diversity distances between obesity groups were compared with pairwise permutational multivariate analysis of variance, based on the vegan package v2.5.7 [47]. The difference in alpha diversity measurements was evaluated with Wilcoxon signed-rank test, implemented in the rstatix v0.7.0 package [57]. In order to evaluate if the obesity groups can be classified based on abundance of bacterial taxa and inferred functional abundances (based on EC, KO, and MetaCyc classification), we used linear discriminant analysis, implemented in in the MASS package v7.3-51.4 [58]. In addition, we repeated linear discriminant analysis using only the 15 most abundant bacterial taxa, in order to evaluate if the dominant bacterial taxa were sufficient for discrimination of the communities, with the obesity status. The results were visualized by plotting the first and second linear discriminants, with the ggplot2 v3.2.1 and the ggpubr v0.4.0 packages [59, 60]. The difference in abundances of bacterial taxa and predicted ECs, KOs, and MetaCyc pathways, between obesity groups was evaluated with a pairwise t-test function, implemented in R v3.6.1 [43]. The correlations between abundances of bacterial taxa were calculated with Spearman’s rank correlation test, included in the Hmisc v4.5.0 package [61]. Resulted correlation matrices were used to construct network plots, using the corrr v0.4.3 package [62]. In addition, the absolute values of correlation coefficients were compared between obesity groups with a pairwise Wilcoxon signed-rank test, implemented in the rstatix v.7.0 package [57]. The influence of 133 recently consumed (from 3-day food records) and 185 habitually consumed (from FFQ) nutrients on beta diversity distances was evaluated with permutational multivariate analysis of variance using a quadratic model [47]. The quadratic model was used as most living organisms, including bacteria have an optimal range of environmental conditions rather than a linear relationship [63-65].

For enrichment analysis, we calculated the mean abundance of each KEGG ortholog for obesity groups and used them as input for MicrobiomeAnalyst (2021-07-01) shotgun data profiling tool, with the default settings [66]. False discovery rate (FDR) *P-*values were adjusted using the Benjamini–Hochberg correction, implemented in rstatix v0.7.0 package [57]. We note that usage of any particular FDR threshold is ambiguous and often varies between microbiome studies; weaker correlations that fail to hold up to *p* adjustment methods often have biological relevance. Premature rejection of associations falling below conservative p-value thresholds may lead to loss of biologically meaningful data. [67-72]. For this reason, statistical results below 0.05 *p*-value threshold were considered to be significant. However, taking into account the difference in opinions and for the readers’ convenience, we report both unadjusted and FDR-adjusted p-values in supplementary data.

## RESULTS

### Lean, overweight and obese individuals can be separated into distinct groups based on their oral and intestinal microbiota

Evaluating beta diversity distances, we observed that salivary microbiota communities of obese and lean individuals were significantly different as measured with Bray-Curtis and Weighted Unifrac distances (**Supplement Table 1**). Based on linear discriminant analysis (non-overlapping confidence ellipses), obesity classes were separated by the abundances of bacterial ASVs (**Fig. 1A**). Obesity groups were also clearly characterized based on abundance of microbial species, genera, families, and orders but weaker based on classes and phyla (**Supplement Figure 1**).

**Figure 1:**
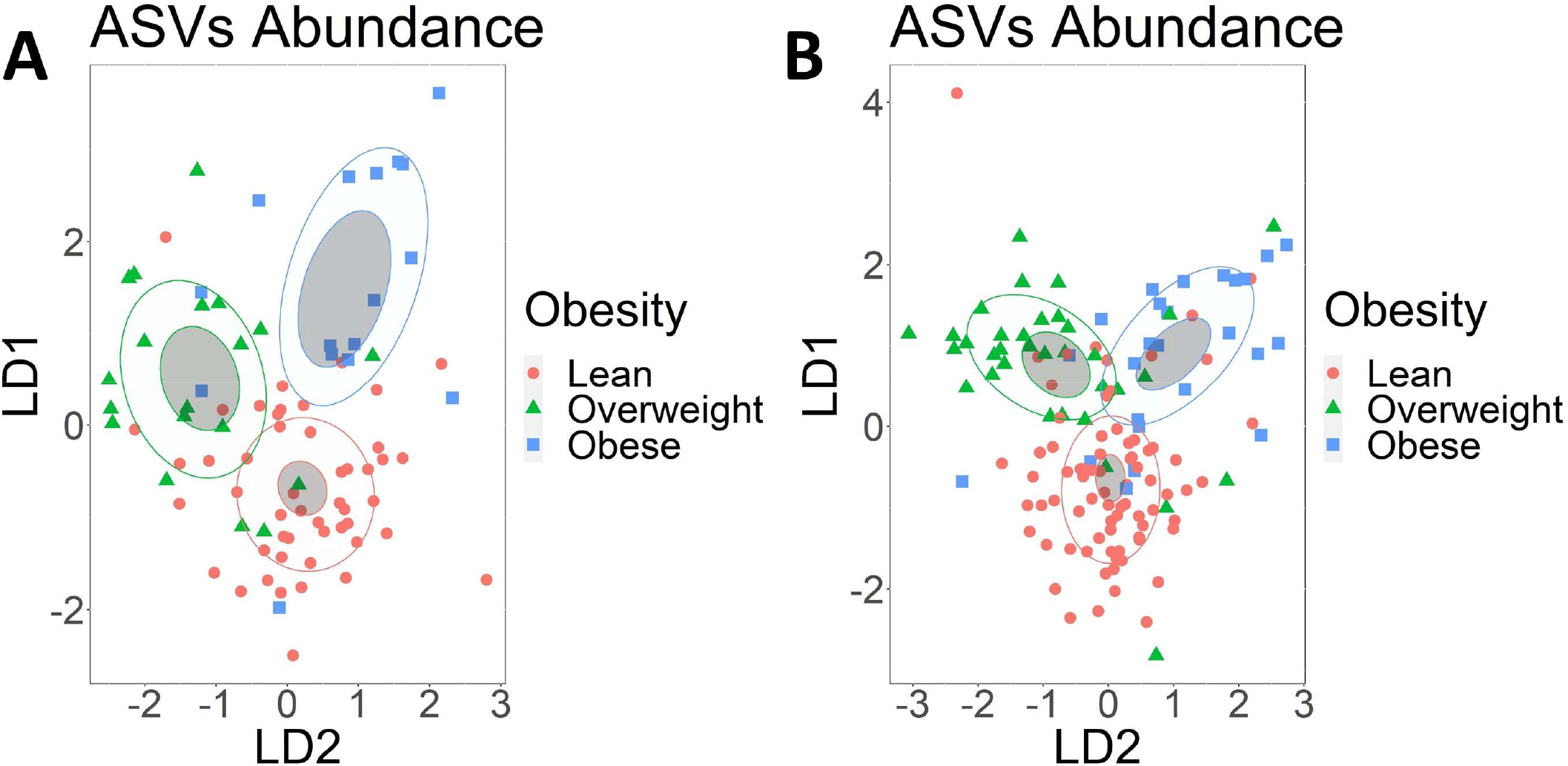
Obesity groups can be discriminated by the abundance of salivary or fecal microbiota. Linear discriminant analysis of A) ASVs identified in salivary samples B) ASVs identified in fecal samples. ASVs with abundance of less than 20 sequences were filtered out. Obesity groups are represented by color, lean group by red, overweight group by green, and obese group by blue. Confidence ellipses are shaded. Normal data ellipses are unfilled and leveled to include 50% of the samples.

In fecal samples, we did not observe a significant difference in beta diversity distances between any of the obesity groups (**Supplement Table 1**). However, based on a linear discriminant analysis, obesity groups could be classified based on abundance of bacterial ASVs (**Fig. 1B**). Obesity groups were also clearly characterized based on abundance of bacterial species, genera, families, and orders but weaker at class and phylum ranks (**Supplemental Figure 2**). We did not observe any significant differences in alpha diversity indices between obesity groups in saliva or feces (**Supplement Table 2**).

### Obesity status influences the abundance of individual bacterial taxa

<u>In saliva,</u> we observed that abundances of Campylobacterota, Firmicutes, and Spirochaetota were significantly different between obesity groups at the phylum rank. Obesity groups were significantly different in the abundances of 5 bacterial classes, 10 orders, 17 families, 33 genera, 52 species, and 409 individual ASVs (**Supplement Table 3A**). Across all taxonomic ranks, obese and lean individuals had the highest number of taxa that were significantly different in their abundances (**Supplement Table 3A**). We evaluated which of the 15 most abundant bacteria taxa were the most influential for defining each of the obesity groups with a linear discriminant analysis. At the genera taxonomic rank, *Campylobacter, Veillonella, Aggregatibacter*, and *Prevotella* defined the obese group (**Fig. 2**). Although lean and overweight groups were not distinct from each other, *Actinomyces* and *Haemophilus* were characteristic for overweight group (**Fig. 2**). Overall, we note that the 15 most abundant bacteria taxa contribute only modestly to discrimination of obesity groups (**Supplemental Fig. 3**).

**Figure 2:**
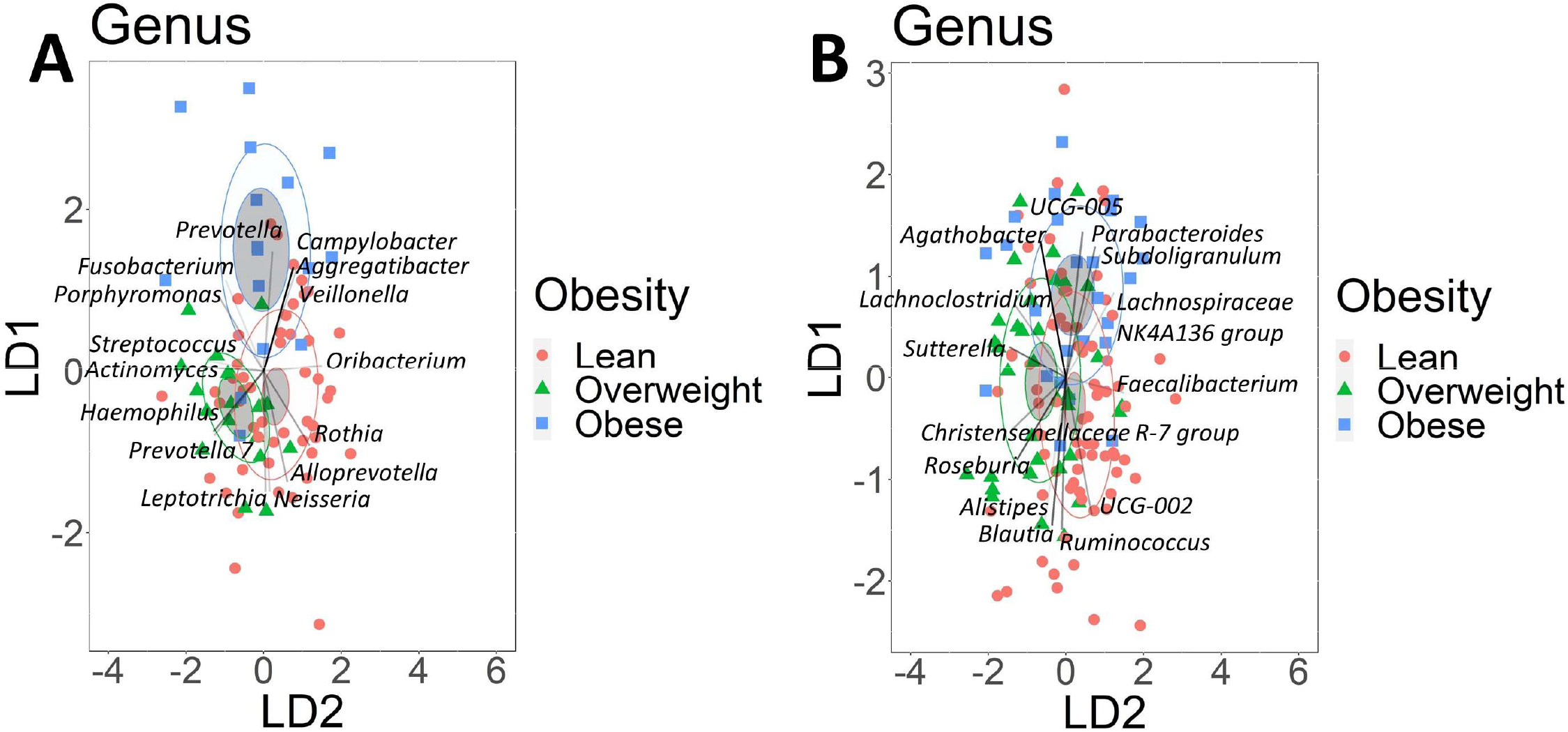
Obese and lean groups can be characterized by the abundance of dominant bacteria genera. Linear discriminant analysis of the 15 most abundant bacterial genera identified in A) Salivary samples B) Fecal Samples. Obesity groups are represented by color, lean group by red, overweight group by green, and obese group by blue. The higher abundance of bacterial genera in the obesity groups is indicated by the direction of the vector rays. The intensity of vector rays’ color corresponds to the strength of the impact. Confidence ellipses are shaded. Normal data ellipses are unfilled and leveled to include 50% of the samples.

<u>In feces,</u> at the phylum rank, only abundance of Fusobacteriota was significantly different between overweight and lean groups. Obesity groups were significantly different in the abundances of 2 bacterial classes, 8 orders, 10 families, 35 genera, 45 species, and 690 individual ASVs (**Supplement Table 3B**). The highest number of significant differences between groups varied with taxonomic rank but was always between lean and one of the overweight/obese groups. Linear discriminant analysis indicated that at the genus taxonomic rank *Agathobacter* and *Parabacteroides* were influential in discriminating obese from lean groups (**Fig. 2**). Although lean and overweight groups were not clearly separated, lean group was primarily characterized by *Blautia* and *Ruminococcus* (**Fig. 2**). Similar to what we observed in salivary samples, the most abundant fecal bacteria taxa were not the most influential variables for discriminating samples based on obesity status (**Supplemental Figure 4**).

### The number of strong correlations between bacterial taxa vary by obesity status

We hypothesized that microbial community inter-relationships, as evidenced by correlations between taxa, would differ by obesity status. We assessed the number of strong correlations (>= |0.7|) between abundances of microbial taxa in saliva and stool samples by obesity group and found evidence for increasing inter-dependence in the setting of obesity (**Fig 3**). Among microbiota genera in saliva, there were 67 strong correlations in the obese group, 32 in the overweight, and only 5 strong correlations in the lean group. The absolute means of correlation coefficients were significantly different between all groups, and this observed pattern remained across all taxonomic ranks (**Supplement Table 4**). We observed a similar pattern in fecal samples, with 52 strong correlations between microbiota genera in the obese group, 20 in the overweight group, and only 8 in the lean group. The absolute values of the correlation coefficients, for abundances of the bacterial taxa were significantly different between all obesity groups. Obese individuals had more strong correlations between bacterial taxa than lean individuals across all phylogenetic ranks except phylum, at which no group had strong inter-bacterial correlations. (**Supplement Table 4**).

**Figure 3:**
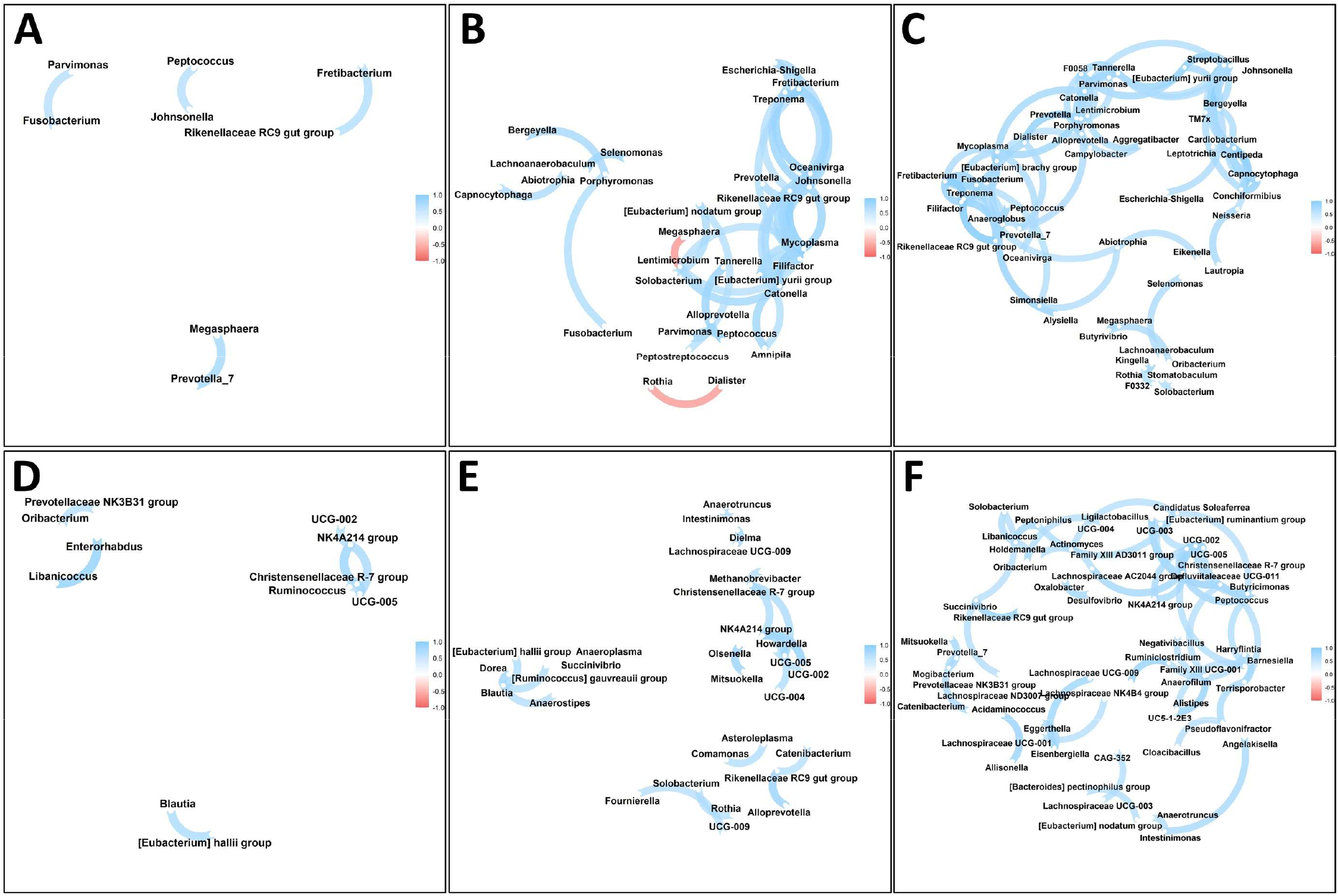
Number of strong connections between bacterial genera increases with the obesity status. Spearman’s rank correlation network between A) Salivary bacterial genera of lean individuals; B) Salivary bacterial genera of overweight individuals; C) Salivary bacterial genera of obese individuals; D) Fecal bacterial genera of lean individuals; E) Fecal bacterial genera of overweight individuals; F) Fecal bacterial genera of obese individuals. For A-C included genera had minimum abundance of 30 sequences and for D-F minimum abundance of 20 sequences.

### Nutritional Factors Influencing Bacterial Communities

We examined the relationships between dietary variables and the overall bacterial community, to identify influential nutrients from recent (3-day food records) and habitual (food frequency questionnaire) consumption. We applied Bray-Curtis, weighted Unifrac, and unweighted Unifrac distances, and assessed both linear and quadratic relationships. For recently-consumed nutrient, xylitol and pectins had significant linear relationships across all 3 methods, while inositol, glucose and omega-3 polyunsaturated fatty acids approached significance for quadratic relationships across all 3 methods (**Supplement Table 5**). For habitually-consumed nutrients, no nutrients displayed consistent linear relationships across all methods, while for quadratic relationships, sorbitol and pinitol, as well as dairy cheese and yogurt were consistently associated (**Supplement Table 6**). In the fecal bacterial community, recently-consumed pectins, folate, and fiber had consistent significant linear relationships, while oxalic acid, formononetin, biochanin A, and the ratio of polyunsaturated to saturated fat had consistent quadratic relationships (**Supplement Table 5**). For habitually-consumed foods, there were consistent linear relationships with cheese and vegetables, in addition to vegetable-derived nutrients (beta carotene, oxalic acid, Vitamin K). Significant quadratic relationships were observed for grains and processed meats, in addition to xylitol, caffeine, sodium and potassium (**Supplement Table 6**).

### Analysis of inferred metabolic pathways reveals enrichment in 2-oxocarboxylic acid metabolism in lean individuals in oral and intestinal microbiota

We hypothesized that functional activity of microbiota, as predicted using PICRUSt2, would differ by obesity status. We assessed differences in inferred function between obesity groups, and found that obesity served as a good classifier for enzyme counts (ECs), KEGG orthologs (KOs), and MetaCyc pathways abundances in saliva (Fig. 4). There were 969 significant differences in ECs, 3,915 in KOs and, 177 significant differences in the abundance of MetaCyc pathways across all groups (**Supplement Table 7**). In all cases, lean and obese individuals had the highest number of differences. 2-oxocarboxylic acid metabolism, terpenoid-quinone biosynthesis, and D-glutamine and D-glutamate metabolism KEGG pathways were enriched in lean individuals but not in obese group (**Supplement Table 8**). The obese group was uniquely enriched in fluorobenzoate, sulfur, and several amino acid metabolic pathways.

**Figure 4:**
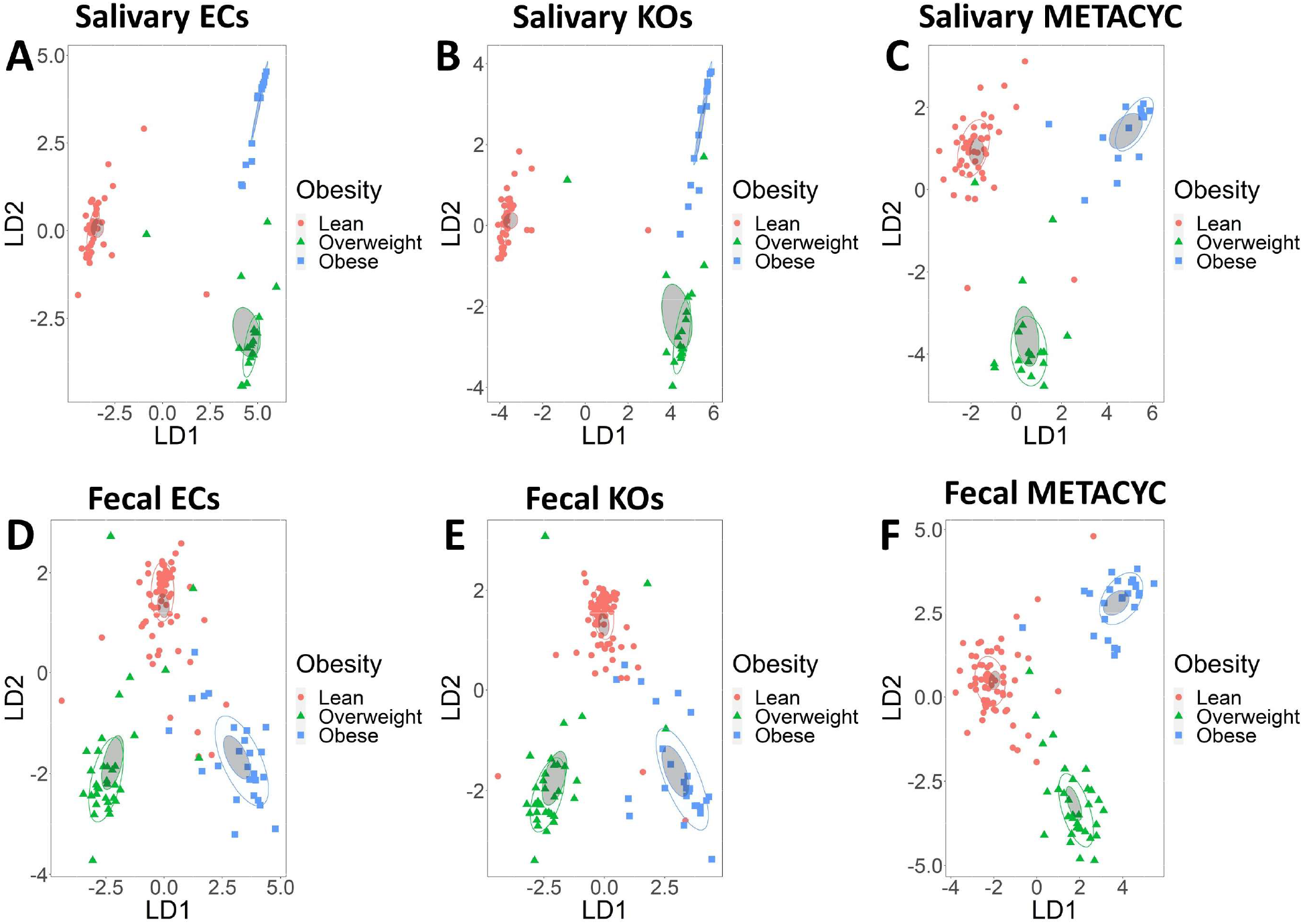
Obesity groups can be discriminated by metabolic potential predicted by PICRUSt2. Linear discriminant analysis of relative abundances of A) ECs inferred from saliva samples B) KOs inferred from saliva samples, C) MetaCyc pathways inferred from saliva samples, D) ECs inferred from fecal samples, E) KOs inferred from fecal samples, F) MetaCyc pathways inferred from fecal samples. Obesity groups are represented by color, lean group by red, overweight group by green, and obese group by blue. Confidence ellipses are shaded. Normal data ellipses are unfilled and leveled to include 50% of the samples.

Similarly, obesity groups could be characterized based on abundance of MetaCyc pathways, KOs, and ECs in fecal samples (Fig. 4). We observed 128 significant differences between the obesity groups in ECs, 391 in KOs, and 19 in MetaCyc pathways (**Supplement Table 7**), spread across lean, overweight and obese groups. The lean group was uniquely enriched in 2-oxocarboxylic acid metabolism, D-glutamine and D-glutamate metabolism, and pentose and glucuronate interconversions, when compared with obese group. The obese group was enriched in C5-branched dibasic acid, lipoic acid, and one-carbon KEGG metabolic pathways (**Supplement Table 8**).

### Abundance of inferred bacterial metabolic enzymes/pathways influences host’s metabolites’ concentrations

We were interested in whether predicted functional activity would associate with measured metabolic activity, as assessed by metabolomic profiling of plasma and stool. We observed high numbers of correlations with predicted saliva microbial activity across all 3 databases (EC: 78,635 with plasma, 82,722 with stool; KO: 249,473 plasma, 263,616 stool; MetaCyc: 15,633 plasma, 17,915 stool). The highest number of correlations was observed with valerate and isoeugenol sulfate in plasma samples and with inosine in stool samples (**Supplement Table 9**). We similarly observed high numbers of correlations between predicted stool microbial activity and metabolites (EC: 92,852 with plasma, 109,830 with stool; KO: 299,557 plasma, 332,789 stool; MetaCyc: 18,179 plasma, 17,728 stool). The highest number of correlations was observed with 1-palmitoyl-GPE and CMPF in plasma samples and steviol in stool samples (**Supplement Table 9**).

## DISCUSSION

Obesity has been linked to alterations in microbiota, however the relative importance of gut and oral microbiota is unclear. We aimed to identify microbial signatures of obesity using both stool and salivary samples in healthy individuals classified as normal weight, overweight or obese based on their BMI. We observed that obesity status was associated with differences in bacterial community composition and shifts in inter-microbial relations that were especially evident in the salivary bacterial community. Although salivary and fecal microbiota were largely impacted by different nutrients, dietary sweeteners were associated with both composition and phylogenetic diversity of both the oral and gut bacterial communities. In addition, samples from obese and lean individuals were enriched in several unique metabolic pathways, inferred activity of which was correlated with plasma and stool metabolite concentrations.

### Obesity influences microbial community composition, especially in saliva

In agreement with published research, we observed that oral bacterial community composition was distinct between lean and obese individuals [24-27]. In our work, we also observed that the difference in salivary bacterial composition between obese and lean individuals extends to phylogenetic diversity measurements. Consistent with previous research, we also observed some differences in gut bacterial communities between obese and lean groups, however in our work the differences were not supported by Bray-Curtis or weighted Unifrac distances [73, 74]. Our results suggest that at the level of the whole community, salivary microbiota composition better reflects the difference in obesity status than fecal microbiota.

With the analysis restricted to the dominant bacterial taxa, we observed a strong influence of *Campylobacter, Aggregatibacter, Veillonella*, and *Prevotella* on characterizing the obese group in salivary samples. Interestingly, all of these bacterial genera have been shown to be correlated not only with obesity but also with oral diseases, especially periodontitis [28-30, 75-77]. Considering the whole bacterial community (abundance >20 reads), we observed that some of the bacteria taxa with lower abundance had a stronger effect on differentiation of the obese group than dominant bacteria, including *Shuttleworthia* at the genus rank and Mycoplasmataceae at the family rank that were also significantly more abundant in the obese group. Previous studies identified a correlation between Mycoplasmataceae and obesity [78, 79]. Although to the best of our knowledge, no previous works associated *Shuttleworthia* with obesity in humans, it was associated with obesity and elevated weight in model organisms [80-82]. In addition, similar to what we observed with the dominant bacteria taxa, *Shuttleworthia* and Mycoplasmataceae are associated with periodontitis [83, 84].

In the fecal samples, the dominant bacterial genera that characterized the obese group were *Agathobacter* and *Parabacteroides. Agathobacter* and *Parabacteroides* were shown to be associated with metabolic disorders in humans and a murine model [85-88]. Similar to what we observed in the saliva samples, several less abundant bacterial taxa that were previously associated with obesity, including *Mitsuokella* and *Neisseria*, at the genus rank and Fusobacteriaceae and Gemellaceae, at the family rank, produced more impact on separation of obese and lean categories than dominant bacterial taxa [73, 89-92]. Proportionally to all identified taxa, more organisms were significantly different in abundance between lean and obese groups in saliva samples, when compared with fecal samples, which might suggest that sampling oral microbiota may be more informative in identifying microbial biomarkers of obesity. Given the relative ease of collection of saliva as compared with stool, this could facilitate increased accessibility for research into the microbial contributors to obesity and cardiometabolic disease; however this remains to be confirmed in independent studies.

### Number of strong correlations between bacterial taxa increases with the obesity status

In saliva samples, bacterial taxa exhibited the highest inter-microbial connectivity (strong correlations >= 0.7) in obese individuals. In the obese group, the highest connectivity was observed for *Fretibacterium* (eight connections), *F0058* (seven connections), *Mycoplasma* (seven connections), and *Tannerella* (seven connections). Several of these genera, including *Fretibacterium, F0058*, and *Tannerella* were shown to be correlated with metabolic disorders [31, 93-96]. In addition, all of the most connected bacterial taxa were associated with periodontitis [83, 94, 97, 98]. In the lean group, the most connected bacteria exhibited less strong connections than in obese group and were *Atopobium* (three connections), *Megasphaera* (two connections), and *Prevotella* 7 (two connections). Abundance of *Atopobium* was shown to be reduced in obese individuals [99]. Previous research indicated that the abundance of *Megasphaera* might increase after anti-obesity treatments [100, 101]. *Prevotella* was shown to be associated with plant rich diet and increase in abundance after antidiabetic treatment, however the genus is very diverse [102-104].

In the fecal samples, the most connected bacterial genera identified in obese group were *Christensenellaceae R7* group (eight connections) and *Ruminococcaceae UCG-005* (five connections). *Christensenellaceae R7* and *Ruminococcaceae UCG-005* were shown to be associated with plasma lipoproteins and triglycerides [105]. *Ruminococcaceae UCG-005* was also shown to be positively correlated with body weight and weight gain in a swine model [106, 107]. In addition, several bacterial taxa previously implicated in metabolic disorders, including *Actinomyces, Ruminiclostridium*, and *Lachnospiraceae* exhibited strong inter-bacterial correlations in the obese but not in the lean group [74, 108-110]. The most connected genus in lean individuals was *Ruminococcaceae NK4A214* (three connections). Previous research identified a negative correlation between *Ruminococcaceae NK4A214* and high fat diet and hypertension [111, 112]. However, *Christensenellaceae R-7* group and *Ruminococcaceae UCG-005* were also among few genera (total three) that had more than one strong correlation in lean individuals.

The impact of the higher degree of microbial interconnectivity observed in obese individuals is unclear but may represent a shift from relative independence of bacterial taxa to a state more reliant on mutualistic relationships. Obesity is often associated with several physiological and environmental conditions that have the potential to act as stressors for the microbial community, including micronutrient deficiency, increased levels of reactive oxygen species, and increase in c-reactive protein concentrations and inflammatory response in the host [113-116]. In accordance with the stress gradient hypothesis, several studies demonstrated that presence of environmental stressors often increases positive facilitation between microbial taxa in the community [117-120]. In addition, it was demonstrated that nutritional stress could increase the number of connections, in a co-occurrence network of the microbiota members [121]. In agreement with these observations, we found that in the obese individuals, almost all of the strong inter-microbial correlations were positive.

### Sweeteners and other nutrients influence compositional and phylogenetic diversity of salivary and fecal bacterial communities

We observed that recently and habitually consumed nutrients influenced bacterial communities. For salivary samples, recently consumed nutrients influenced bacterial community more than habitually consumed nutrients, for both compositional and phylogenetic beta diversity distances. Sugars and sugar alcohols, especially xylitol, mannitol, sorbitol, and pectin were especially influential factors impacting the bacterial community, based on compositional and phylogenetic diversity measurements. Interestingly, all of the listed compounds with the exception of pectin are used as sweeteners [122, 123]. Although the effect of sweeteners on gut microbiota was extensively shown in humans and animal models, the studies on oral bacteria community are limited [124, 125]. To the best of our knowledge, this work is the first report on the correlation between dietary sweeteners and phylogenetic diversity of the human’s salivary bacterial community.

Fecal microbiota community was consistently more influenced by habitual nutrient consumption than recently consumed nutrients, which might suggest a more stable microbial community. Similar to the saliva samples, consumption of xylitol and pectin influenced compositional and phylogenetic diversity of fecal microbiota. Consumption of sweeteners, including xylitol was reported to influence intestinal bacterial community composition [124]. Pectin consumption was also shown to be correlated with compositional changes in the intestinal microbiota [126, 127]. In our study, compositional and phylogenetic measurements of the fecal microbiota were also consistently influenced by consumption of vegetables and plant-derived compounds including fiber, oxalic acid, formononetin, and daidzein. Consumption of fiber, formononetin, and daidzein was show to have microbiota-mediated beneficial effects on host’s metabolic health [128, 129] In addition, habitual consumption of cholesterol and fatty acids also produced a significant effect on compositional and phylogenetic diversity distances of the fecal microbiota in our study.

### Bacterial communities of obesity groups are associated with enrichment in predicted metabolic pathways, which are correlated with host’s metabolite concentrations

In both saliva and fecal samples, microbiota of the lean individuals were enriched in 2-oxocarboxylic acid metabolism and D-glutamine and D-glutamate metabolism, based on functional prediction. 2-Oxocarboxylic acid metabolism is involved in ornithine and lysine biosynthesis, supplementation of which were shown to have a potential for improving metabolic health [130-132]. D-Glutamine concentrations were shown to be decreased in obese individuals and glutamine supplementation may alleviate obesity symptoms [133, 134]. Metabolic pathways enriched in the microbiota of obese individuals included one-carbon metabolism, which was previously shown to contribute to the development of obesity[135]. In addition, steatosis was shown to be associated with one carbon metabolism’s gene expression [136]. Enrichment in other pathways such as lipoic acid metabolism and degradation of valine, leucine, and isoleucine might be a response to increase in oxidative stress and branched-chain amino acids concentrations, often associated with obesity [137-139].

Multiple host’s metabolites were significantly correlated with abundance of KOs involved in enriched pathways. For example, the abundance of KOs, predicted in salivary samples and involved in 2-oxocarboxylic acid metabolism influenced the concentration of 435 plasma and 326 stool metabolites. Alpha-ketobutyrate was shown to be a biomarker of insulin resistance and glucose intolerance and in our study exhibited a negative correlation with more than half of the 2-oxocarboxylic acid metabolism pathway’s KOs, predicted from saliva samples [140, 141]. In addition, KOs involved in 2-oxocarboxylic acid metabolism were correlated with adenosine and steviol in stool samples, both of which were shown to be beneficial for patients with metabolic disorders [142, 143].

Our study had considerable strengths, including availability of salivary and fecal microbial profiling, in addition to metabolic phenotyping, in a robust sample size. There were also some limitations inherent in all microbiome projects that are based on 16S rRNA sequencing. Namely, the necessity of choosing a specific segment of the gene, sequence filtering methods, reference database for taxonomic identification, and even normalization methods are all know to cause a degree of bias between studies. In addition, results presented in this study are largely based on relative abundances of the identified microbial taxa and therefore might not be interpreted as causative. Therefore, future studies would be necessary to demonstrate the directions of interactions between the host and its oral and intestinal microbiota.

## CONCLUSIONS

In this study we identified differences in salivary and fecal symbiotic bacterial communities based on obesity status, in a population of otherwise healthy individuals. Our results suggest that inter-correlations between bacterial taxa are altered in the setting of obesity and suggest distinct differences in community dynamics at increasing levels of obesity. Consideration of microbial community correlation structure might be more informative than measurement of relative abundances of bacteria taxa or diversity measurements alone. In addition, across multiple comparisons, salivary microbiota provided a more distinct pattern of differentiation between obese and lean individuals, than fecal microbiota. Previous studies have primarily focused on analysis of gut microbiota in obesity, however our data suggest that sampling oral microbiota might be a better choice in search of the bacterial biomarkers associated with obesity.

## Acknowledgements

This work was supported by funding from NIH R01 HL142856, and the Layton Family Fund.

## Supplementary Figures’ Legends

**Supplement figure 1: Obesity groups could be characterized based on abundance of salivary bacterial taxa, especially at lower taxonomic ranks**. Linear discriminant analysis of A) bacterial species, B) bacterial families, C) bacterial families, D) bacterial orders, E) bacterial classes F) bacterial phyla. Taxa with abundance of less than 20 sequences were filtered out. Obesity groups are represented by color, lean group by red, overweight group by green, and obese group by blue. Confidence ellipses are shaded. Normal data ellipses are unfilled and leveled to include 50% of the samples.

**Supplement figure 2: Obesity groups could be characterized based on abundance of fecal bacterial taxa, especially at lower taxonomic ranks**. Linear discriminant analysis of A) bacterial species, B) bacterial genera, C) bacterial families, D) bacterial orders, E) bacterial classes F) bacterial phyla. Taxa with abundance of less than 20 sequences were filtered out. Obesity groups are represented by color, lean group by red, overweight group by green, and obese group by blue. Confidence ellipses are shaded. Normal data ellipses are unfilled and leveled to include 50% of the samples.

**Supplement figure 3: Lean and obese groups can be characterized by the abundance of dominant salivary bacterial taxa, across most taxonomic ranks**. The results also indicate that the most abundant bacteria taxa are not the most influential for characterization of obesity groups. Linear discriminant analysis of 15 most abundant salivary bacterial A) Species, B) Families, C) Orders, D) Classes. Obesity groups are represented by color, lean group by red, overweight group by green, and obese group by blue. The intensity of vector rays’ color corresponds to the strength of the impact. Confidence ellipses are shaded. Normal data ellipses are unfilled and leveled to include 50% of the samples.

**Supplement figure 4: Lean and obese groups can be characterized by the abundance of dominant fecal bacterial taxa across most taxonomic ranks**. The results also indicate that most abundant bacteria taxa are not the most influential for characterization of obesity groups. Linear discriminant analysis of 15 most abundant fecal bacteria A) Species, B) Families, C) Orders, D) Classes. Obesity groups are represented by color, lean group by red, overweight group by green, and obese group by blue. The intensity of vector rays’ color corresponds to the strength of the impact. Confidence ellipses are shaded. Normal data ellipses are unfilled and leveled to include 50% of the samples.

## Supplementary Tables’ Legends

**Supplement Table 1: Salivary but not fecal samples exhibited a significant difference in compositional and phylogenetic distances between lean and obese groups**. Pairwise comparisons of Bray-Curtis, Weighted and Unweighted Unifrac distances between obesity groups in salivary and fecal samples.

**Supplement Table 2: Obesity groups did not exhibit a significant variation in alpha diversity indices of salivary and fecal microbial communities**. Pairwise comparisons of Shannon and Gini-Simpson indices’, as well as species richness values between obesity groups in salivary and fecal samples.

**Supplement Table 3: More salivary bacterial taxa exhibited a significant difference in their abundances between obesity group than fecal bacterial taxa**. Pairwise comparisons of microbiota taxa abundances between obesity groups in A) salivary samples and B) fecal samples.

**Supplement Table 4: The number of strong correlations between bacterial taxa in salivary and fecal samples increased with the increase of the obesity status of the host**. Pairwise comparisons of absolute values for inter-bacterial correlation coefficients between obesity groups.

**Supplement Table 5: More of the recently consumed nutrients produce a significant effect on compositional and phylogenetic diversity distances of salivary samples than on fecal samples**. The impact of changes in recently consumed nutrients on Bray-Curtis and Weighted and Unwheighted Unifrac distances of salivary and fecal samples. Sq stands for quadratic effect of the nutrient on beta diversity distances.

**Supplement Table 6: More of the habitually consumed nutrients produce a significant effect on compositional and phylogenetic diversity distances of fecal samples than on salivary samples**. The impact of changes in habitually consumed nutrients on Bray-Curtis and Weighted and Unwheighted Unifrac distances of salivary and fecal samples. Sq stands for quadratic effect of the nutrient on beta diversity distances.

**Supplement Table 7: Predicted metabolic potential of bacterial community varies between obesity groups**. Pairwise comparisons of predicted bacterial enzymes, Kegg orthologs, and MetaCyc abundances between obesity groups in A) salivary samples and B) fecal samples. ECs, KOs, and MetaCyc pathways that were not significantly different (*p* >0.05) between obesity groups were excluded from the table.

**Supplement table 8: Some of the predicted bacterial metabolic pathways are uniquely enriched in obesity groups**. Pairwise comparisons of uniquely enriched bacterial metabolic pathways identified from salivary and fecal samples between obesity groups.

**Supplement table 9: Predicted bacterial metabolic potential influences plasma and stool metabolites concentrations**. The effect of A) ECs identified from salivary samples on plasma metabolites concentrations, B) ECs identified from salivary samples on stool metabolites concentrations, C) ECs identified from fecal samples on plasma metabolites concentrations, D) ECs identified from fecal samples on stool metabolites concentrations, E) KOs identified from salivary samples on plasma metabolites concentrations, F) KOs identified from salivary samples on stool metabolites concentrations, G) KOs identified from fecal samples on plasma metabolites concentrations, H) KOs identified from fecal samples on stool metabolites concentrations, I) MetaCyc pathways identified from salivary samples on plasma metabolites concentrations, J) MetaCyc pathways identified from salivary samples on stool metabolites concentrations, K) MetaCyc pathways identified from fecal samples on plasma metabolites concentrations, L) MetaCyc pathways identified from fecal samples on stool metabolites concentrations.

